# Phylogenetic and functional compositional shifts associated with selective logging and forest regrowth in tropical rainforests of the Congo Basin

**DOI:** 10.1101/2024.10.31.621313

**Authors:** Jonas Depecker, Justin A. Asimonyio, Yves Hatangi, Olivier Honnay, Steven B. Janssens, Jean-Léon Kambale, Filip Vandelook, Angelino Carta

**Author notes:** Corresponding author: Jonas Depecker.

## Abstract

**Background and aims:** Tropical rainforests constitute a globally important biome severely threatened by anthropogenic activities. Accounting for the phylogenetic and functional dimensions can provide insights into how anthropogenic activities affect tree species community assembly. Here, we aimed to assess the effects of selective logging and forest regrowth on the tree composition in the Yangambi region (Democratic Republic of the Congo), as compared to reference undisturbed old-growth forest by incorporating phylogenetic and functional information.

**Methods:** We measured the phylogenetic and functional alpha-diversity and dissimilarity of undisturbed old-growth, disturbed old-growth, and regrowth rainforests by sampling species abundances and key traits related to usefulness (wood density), vegetative (specific leaf area), and reproductive (fruit type) functions. Then, we evaluated the association between phylogenetic and functional dissimilarity to unravel potential drivers underlying changes in the community composition.

**Key results:** Phylogenetic and functional dissimilarities resulted in a consistent compositional separation of the disturbed old-growth forests and regrowth forests from the undisturbed old-growth forests, with this separation clearly associated with key functional traits.

Compared to undisturbed old-growth forests, disturbed old-growth forests subjected to selective logging did not diverge in terms of both phylogenetic and functional diversity and structure. However, regrowth forests displayed increased levels of phylogenetic diversity and comparable functional diversity as undisturbed old-growth forests.

**Conclusions:** Selective logging and forest regrowth have not led to an overdispersion or clustering of phylogenetic lineages nor functional traits in the tropical rainforests in the Congo Basin, rather, these anthropogenic activities brought an altered phylogenetic and functional community composition which may have serious implications for the stability of these ecosystems.

## Introduction

The Afrotropical rainforests stand out as one of the most crucial biomes worldwide, renowned for their importance in the conservation of floristic diversity and the delivery of essential ecosystem services and functions (Atangana *et al*., 2013; Grantham *et al*., 2020). Over 30,000 vascular plant species are known for this region, with most angiosperm lineages represented (Ramírez-Barahona *et al*., 2020; Raven *et al*., 2020; Carta *et al*., 2022). However, compared to the tropical rainforests in the Neotropical region, plant diversity in the Afrotropical region is lower, which can be attributed to differences in geological and climatic history, as well as the significant under-sampling of this region (Sosef *et al*., 2017; Raven *et al*., 2020). In terms of ecosystem functions, the Afrotropical rainforests play a key role in gross primary production and, more importantly, in carbon storage (Hubau *et al*., 2020; Harris *et al*., 2021). The rainforests of the Congo Basin alone, accounting for 89 % of the Afrotropical rainforests, sequester 0.61 Gt of atmospheric CO_2_ per year, six times the amount sequestered by the Amazon region of which the carbon sink has been declining since the 1990s (Malhi *et al*., 2013; Hubau *et al*., 2020; Harris *et al.,* 2021). Despite the importance of these rainforests for biodiversity conservation and for providing ecosystem services, anthropogenic activities are increasingly threatening their existence (Tyukavina *et al*., 2018). According to the FAO, about four million ha of African tropical rainforest is lost annually, at a rate twice as fast as elsewhere in the world (FAO & UNEP, 2020).

Besides the conspicuous deforestation, a much less detectable anthropogenic threat to the tropical rainforest, but potentially more detrimental based on the extensive spatial scale on which it occurs, is the degradation of tropical rainforests (Barlow *et al*., 2016). Forest degradation is the anthropogenic disturbance within the forest, under a more or less intact canopy layer, and entails a plethora of anthropogenic activities of which selective logging is one of the main activities, especially in the Afrotropics where selective logging is mainly driven by the production of charcoal and extraction of construction wood (Sasaki & Putz, 2009; Simula, 2009; Hosonuma *et al*., 2012). Because of the large spatial scale at which forest degradation occurs, most of the Afrotropical rainforest is nowadays affected by logging with one-third considered to be heavily degraded (Gibson *et al*., 2011; Döbert *et al*., 2017; Shapiro *et al*., 2021). Currently, the annual rate of forest degradation in the Congo Basin alone is estimated at nearly 1.5 million ha (Shapiro *et al*., 2021).

Although tropical deforestation rates remain alarmingly high, the regrowth of abandoned tropical rainforest has become an important process as well. A substantial part of tropical rainforests consists of regrowth forests, and this proportion is expected to increase. Many African countries are committed to restore deforested lands to their natural forest state, including through initiatives like the African Forest Landscape Restoration Initiative (AFR100) (Ernst *et al*., 2013; Berenguer *et al*., 2018; CIFOR, 2022; AFR100, 2024). As such, nearly one million forest ha have been actively or passively restored in Sub-Sahara Africa between 2000 and 2019 (FAO & UNEP, 2020). Studies from outside the Congo Basin suggest that species richness and diversity in regrowth forests tend to recover relatively quickly. Contrarily, it takes longer for the species composition of these forests to reach pre-deforestation levels (Hayward *et al*., 2021; Oberleitner *et al*., 2021; Poorter *et al*., 2021). However, further research is required to determine if these patterns also apply to the rainforests of the Congo Basin.

The effects of forest degradation and forest regrowth on biodiversity are commonly assessed with traditional measures of taxonomic richness, diversity, or species composition analyses (Döbert *et al*., 2017). However, as already recognised by Darwin (1859), species are not independent entities in terms of their shared evolutionary history and functional diversity (Pausas & Verdú, 2010; Garamszegi, 2014). Taxonomic assessments lack a considerable part of biological diversity and its related processes (Chave *et al*., 2007; Hernández□Ordóñez *et al*., 2019) and should also be analysed in a phylogenetic and functional dimension (Faith, 1992; Swenson, 2011; Pollock *et al*., 2017). The phylogenetic dimension accounts for the variation in ancestral relatedness, including whether taxa are rooted more at the base (“basal”) or at the tips (“terminal”) within a given phylogenetic tree, providing insight into the shared evolutionary history and divergence of taxa (Garamszegi, 2014). The functional dimension considers variation in functional trait values and provides a perspective of the ecological niche of taxa, and the ecosystem functioning of the communities (Döbert *et al*., 2017; Tucker *et al*., 2017; Castillo-Campos *et al*., 2024; Bañares-de-Dios *et al*., 2024). Conceptually, phylogenetic and functional dimensions could be closely related, with the underlying assumption that more closely related species tend to share more similar functional traits (Pavoine *et al*, 2011; Bañares-de-Dio *et al*., 2024). However, studies on the relation between phylogenetic and functional diversity yield inconsistent results and, the underlying assumption has been challenged numerous times (Tucker *et al*., 2018; and references within). Indeed, a mismatch between the different diversity metrics may exist, in which the phylogenetic diversity is decoupled from the functional diversity, or *vice versa* (Devictor *et al*., 2010). Furthermore, anthropogenic processes may act differently on phylogenetic diversity than on functional diversity (Devictor *et al*., 2010). An integrative approach that measures taxonomic, phylogenetic, and functional diversity together is essential for a better understanding of how forest degradation and regrowth affect the structure, composition, and dynamics of tropical rainforest tree communities. This comprehensive assessment is crucial for developing effective conservation strategies (Devictor *et al*., 2010; Döbert *et al*., 2017).

Although the importance of phylogenetic and functional diversity is well recognised, studies that examine the effects of forest degradation and regrowth by integrating both the phylogenetic and functional dimensions remain limited, especially in the tropics. From the few available studies in Asia (Ding *et al*., 2012; Toyama *et al*., 2015; Ding *et al*., 2019; Cao *et al*., 2021) and South America (Baraloto *et al*., 2012; Carreño□Rocabado *et al*., 2012; Ribeiro *et al*., 2016; Rocha-Santos *et al*., 2023), there seems to be a common pattern of decreasing diversity with increasing disturbance, accompanied by a significant changes in both phylogenetic and functional composition (Mouillot *et al*., 2013; Döbert *et al*., 2017; Comita *et al*., 2018). In the Afrotropics, and particularly in the Congo Basin, such studies are, to the best of our knowledge, lacking. In terms of forest regrowth, there seems to be a trend in fast recovery of taxonomic diversity, yet not at the level of phylogenetic and functional composition (Mo *et al*., 2013; Mahayani *et al*., 2020; Mahayani *et al*., 2022). Similarly, only few studies have been conducted on this topic in Asia (Mo *et al*., 2013) and South America (Whitfeld *et al*., 2012; Mahayani *et al*., 2020; Mahayani *et al*., 2022). However, research on this topic in the Congo Basin remains scarce, with some exceptions (e.g. Makelele *et al*., 2021).

Here, we aim to fill this knowledge gap by leveraging previously collected data by Depecker *et al*. (2022) to assess the impact of forest degradation through selective logging and of forest regrowth on the phylogenetic and functional diversity and composition of tropical rainforest, as compared to reference undisturbed old-growth forest in the Congo Basin. Depecker *et al*. (2022) established 125 sampling quadrats across 25 rainforest inventory plots in the Yangambi region in the Democratic Republic of the Congo (DR Congo) showing that, on the one hand, logged forests were clearly associated with a reduced number of tree individuals with a lower wood density and an altered taxonomic community composition. On the other hand, regrowth forests showed a relatively fast recovery of taxonomic diversity, but not of taxonomic tree community composition and forest structure after agricultural abandonment. Based on these outcomes, we expect: (i) to observe a significant overlap in phylogenetic and functional dissimilarities, since both dimensions are often conceptually related and selective logging is removing only a few species with high wood density, (ii) that terminal phylogenetic diversity will be more affected rather than the basal phylogenetic diversity, since only a selected number of tree individuals and species are removed, while the original diversity in plant families continue to be represented after selective logging, and (iii) to find a significantly altered phylogenetic community composition in regrowth forest, driven by species with a low wood density.

## Material and methods

### Study area and vegetation sampling

The Yangambi region (northeastern DR Congo), is located approximately 100 km west of the major city Kisangani. Yangambi serves as a prime example of tropical rainforest landscapes within the Congo Basin, *i.e.* a mosaic of land tenures. The landscape is primarily composed of the Yangambi Man and Biosphere Reserve, the Ngazi Forest Reserve, a timber logging concession, and customary land (van Vliet *et al*., 2018; van Vliet *et al*., 2019). Despite the incorporation of the Yangambi reserve in the Man and Biosphere framework in 1979, no management plan has been implemented up to date. Moreover, the borders of the reserve are still heavily contested (Koy *et al*., 2019; van Vliet *et al*., 2023). In an area spanning approximately 50-by-20 km within this landscape, Depecker *et al*. (2022) established 25 forest inventory plots of 125 m x 125 m across three different forest categories. Classification of the forests was done combining historical land-use maps and contemporary information. As such, (i) plots on historical agricultural and which have been overgrown since abandoned somewhere between 40 and 60 years ago were classified as regrowth forest (R), (ii) plots on historical old-growth forest land with clear indications of small-scale selective logging were classified as disturbed old-growth forest (DO), and (iii) plots on historical old-growth forest land without signs of selective logging were classified as undisturbed old-growth forest (UO).

### Data collection

Vegetation sampling was conducted by Depecker *et al*. (2022) within the 25 established forest inventory plots, using five quadrats of 25 m x 25 m per plot, and then later summed into one value per plot for the further analyses. A total of 7,375 trees with a diameter at breast height (DBH) greater or equal than 5 cm was reported, belonging to 211 taxa. Of these, 167 were identified to the species level, totalling 6,953 trees. Additionally, this data was supplemented with estimates of specific leaf area (SLA) and wood density (WD), obtained from measurements on herbarium specimens and compiled from literature, respectively, for the taxa identified at species level. In this study, the dataset of Depecker *et al*. (2022) was complemented with information on fruit type (FT; achene, berry, capsule, drupe, follicle, legume, and nut) for each taxon identified at species level gathered from literature (Orwa *et al*., 2009; Mouly *et al*., 2014; Kattge *et al*., 2020; Bbidjo *et al*., 2024; Hyde *et al*., 2024; Sosef *et al*., 2024)

### Phylogeny construction

Prior to the construction of the phylogenetic tree, a taxonomic homogenisation of all species names against the World Checklist of Vascular Plants was conducted using the *wcvp_match_names* function in the rWCVP package (Brown *et al*., 2023). The phylogeny of the 167 species was then reconstructed using the U.PhyloMaker package (Jin & Qian, 2023), and based on the “GBOTB.extended.WP” tree (Jin & Qian, 2022) which encompasses over 70,000 species of vascular plants. Taxa in our study that were absent from the Jin and Qian (2022) mega-tree were appended to their closest relative in the mega-tree using the *bind.relative* function. Assessment of the closest relative was based on phylogenetic systematic literature (see supplementary material; Table S1) including another large-scale angiosperm species phylogeny (Janssens *et al*., 2020).

### Data analysis

#### Phylogenetic diversity

To assess the phylogenetic diversity in the tree communities, a range of metrics was calculated based on the previously generated phylogenetic tree and accounting for the number of individuals per species. Three common metrics of phylogenetic diversity were used, each emphasising different depths of evolutionary history across a phylogeny (Faith, 1992; Tucker *et al*., 2017; Qian *et al*., 2017): Faith’s phylogenetic diversity (PD), mean pairwise distance (MPD), and mean nearest taxon distance (MNTD). PD is measured as the sum of all branch lengths in the phylogenetic tree and is considered a richness metric representing the quantity of phylogenetic history present in an assemblage. MPD and MNTD on the other hand, are both divergence metrics based on pairwise cophenetic distances. Whereas the former considers all pairwise distances between taxa in an assemblage and is more sensitive to the basal structure of the phylogenetic tree, the latter considers only the nearest distances between taxa and is more sensitive towards the terminal structure of the phylogenetic tree (Faith, 1992; Tucker *et al*., 2017). It should be noted that basal and terminal in this context should be interpreted relative to the phylogenetic tree used in this study and not *per se* to the whole angiosperm phylogeny.

Since phylogenetic diversity (PD) has a strong positive correlation with species richness (Mazel et al., 2016), the standard effect size (SES) of the aforementioned phylogenetic metrics was calculated to compare plots with different species richness levels. This was done by standardising the metrics using the “taxa.labels” null model, which maintains constant species richness while randomising phylogenetic relationships across 999 iterations (Kembel et al., 2010; Mazel et al., 2016).The standardised metrics quantify the relative overdispersion (excess) or clustering (deficit) in phylogenetic diversity for a given set of species relatively to the species pool. Consequently, positive values of these metrics represent a relative overdispersion of species, whereas negative values of these metrics represent a relative clustering of species (Mazel *et al*., 2016; Gioria *et al*., 2023). All phylogenetic diversity metrics were estimated using the picante package (Kembel *et al*., 2010) and visualised using the ggplot2 package in R (Wickham, 2016; R Core Team, 2024).

Differences in phylogenetic diversity among the three different forest categories were afterwards assessed with generalised linear mixed models (GLMM) with a Gaussian error distribution, selected after inspection of the distribution of the residuals. In the models, the forest categories acted as fixed factors and the spatial cluster of the forest inventory plots as a random factor using the lme4 (Bates *et al*., 2015) and multcomp (Hothorn *et al*., 2008) packages in R (R Core Team, 2024). The spatial clustering of the forest inventory plots was represented by seven spatial groups which were identified by a hierarchical clustering analysis based on the geographical distance between the plots.

#### Functional diversity

Before estimating the functional diversity, the non-continuous fruit type data was transformed into continuous data based on the multidimensional functional spaces framework as outlined by Maire *et al*. (2015). Firstly, the functional dissimilarity is measured between species, which was done using the Gower’s distance (Gower, 1971) using *vegdist* function in the vegan package (Oksanen *et al*., 2022). Secondly, based on the functional dissimilarity matrix the multidimensional functional spaces were computed using a principal coordinates analysis with the *pcoa* function in the ape package (Paradis & Schliep, 2019). Finally, the three first dimensions with eigenvalues greater than one and cumulative relative eigenvalues of approximately 80 % were selected and used as input to calculate the functional diversity indices (FT1, FT2, and FT3). Subsequently, using all continuous functional traits, a range of metrics was calculated representing the most important dimensions of functional diversity (Mouchet *et al*., 2010). The computed functional diversity metrics included: functional richness (FRic) which represents the total amount of functional space filled by the species in a community; functional divergence (FDiv) which represents how abundance (number of individuals) is spread along the different traits; functional evenness (FEve) which describes the regularity of the distribution of species (and their abundances) in trait space; functional dispersion (FDis), reflects changes in the abundance-weighted deviation of species trait values from the centre of the functional space; and Rao’s quadratic entropy (FRaoQ) assesses the multi-dimensional divergence in trait space and is the abundance-weighted variance of the trait dissimilarities between all species pairs (Grenié & Gruson, 2023). All functional diversity metrics were estimated using the fundiversity package (Grenié & Gruson, 2023) on standardised trait data and visualised using the ggplot2 package (Wickham, 2016). Testing for differences in functional diversity metrics between the three forest categories was done using generalised linear mixed models following the same procedure as for the phylogenetic diversity metrics.

In order to investigate the functional structure of the communities in the different forest categories, standardised effect sizes of the abovementioned functional diversity metrics were calculated following the methods outlined by Mastrogianni *et al*. (2021). First, 999 random community data matrices were generated from the original species pool with the richness null model using the *randomizeMatrix* function in the picante package (Kembel *et al*., 2010). Subsequently, all functional diversity metrics were computed for the randomised communities. The SES values were then calculated by subtracting the mean value of the functional diversity metric under the null distribution from the observed value of the diversity metric, and then divided by the standard deviation of the functional diversity metric under the null distribution. In analogy with the SES values of the phylogenetic diversity metrics, positive values of SES functional diversity metrics represent functional overdispersion, whereas negative values represent functional clustering. Finally, following Ortega-Martínez *et al*. (2020), statistical significance at P < 0.05 was inferred when the SES value fell outside the range of –1.96 to 1.96.

#### Phylogenetic-functional community framework

First, the correlation between the phylogenetic and functional diversity metrics was assessed with *cor.test.* Subsequently, a phylogenetic-functional community framework was constructed through a four-step process. Firstly, the phylogenetic and functional beta-diversity were calculated based on the phylogenetic and trait distances among species, respectively, and taking into account the abundance per species. This was done using both the *comdist* and *comdistnt* functions in the picante package (Kembel *et al*., 2010), which measure the equivalents of MPD and MNTD, respectively, for among-communities. The correlation between the phylogenetic and functional beta-diversity, based both on MPD and MNTD, was afterwards tested with multiple mantel tests with 999 permutations. Secondly, both phylogenetic and functional beta-diversity across the forest categories were visualised using separate non-multidimensional scaling (NMDS) ordinations. The ordinations were performed with the *metaMDS* function in the vegan package (Oksanen *et al*., 2022), and for each ordination, a two-axis solution provided a stress level below the 0.20 threshold, indicating a good representation in reduced dimensions. Subsequently, a permutational multivariate analysis of variance with 999 permutations was conducted to test for differences in beta-diversity among forest categories with the *adonis* function in the vegan package (Oksanen *et al*., 2022), followed by pairwise comparisons of the forest categories with 999 permutations and Bonferroni P-value correction with the *pairwise.adonis.function* in the pairwiseAdonis package (Arbizu, 2017). Thirdly, to explore which phylogenetic lineages are possibly driving the observed patterns in beta-diversity, the main clades (monocots, magnoliids, asterids, and rosids) were fitted onto the ordination results with the *envfit* function in the vegan package (Oksanen *et al*., 2022) with 999 permutations. Finally, to explore which traits are possibly driving the observed patterns in beta-diversity, the functional traits were fitted onto the ordination results following the same approach as for the phylogenetic lineages, together with the previously estimated phylogenetic and functional diversity metrics. For this last analysis, the community-weighted means were used for FT, SLA, and WD, whereas for DBH the frequency per quartile was used (DBH1, DBH2, DBH3, and DBH4).

## Results

### Phylogenetic diversity

Overall, the disturbed old-growth forests exhibited a relatively higher basal phylogenetic diversity as compared to undisturbed old-growth forests. Specifically, MPD was 3.51 % higher in DO than in UO (p < 0.01; Figure 2). For PD and MNTD, no significant differences between DO and UO were found. SES values tended to be more positive for MPD and more negative for MNTD in DO than in UO, yet very few of the plots showed significant clustering or overdispersion when compared with the null model.

**Figure 1:**
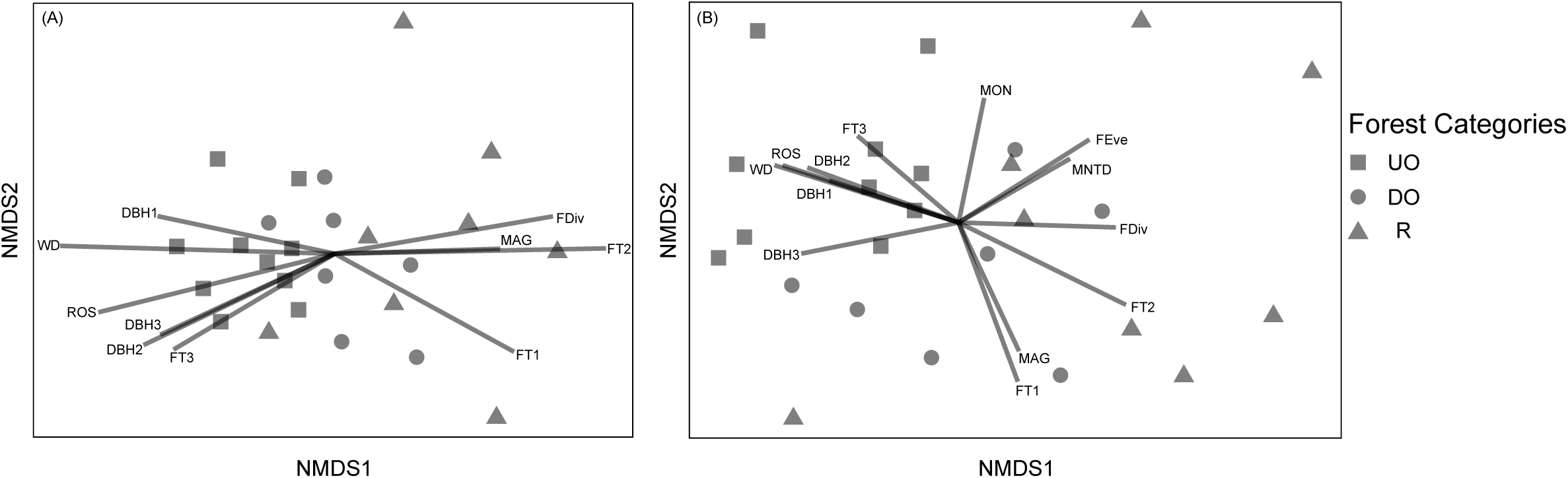
Phylogram depicting the shared evolutionary history of the 167 taxa included in the study, including information on fruit type (FT), specific leaf area (SLA), and wood density (WD). Branches are coloured following APG clades monocots, magnoliids, asterids, and rosids.

**Figure 2:**
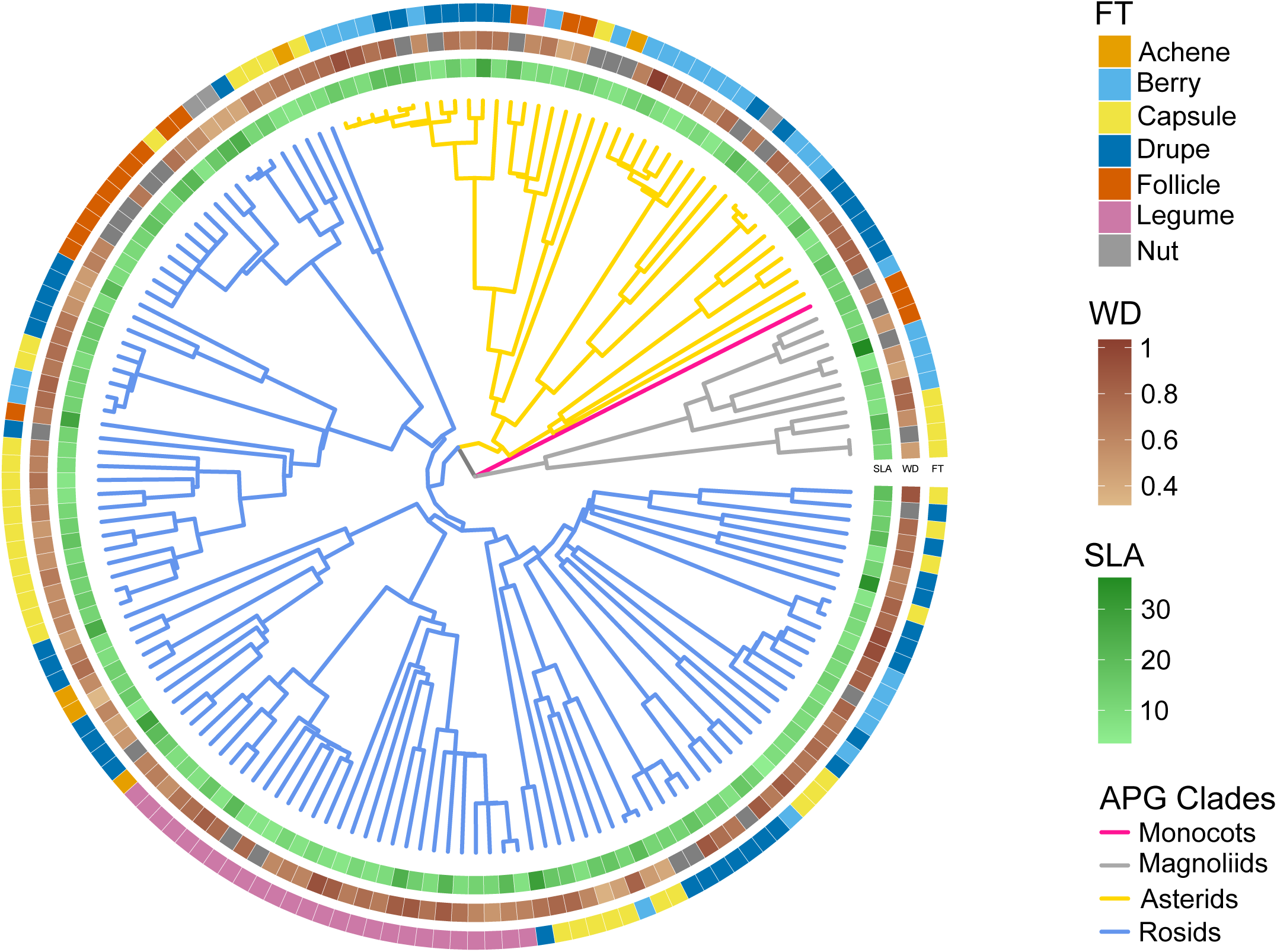
Comparisons of the observed (A) Faith’s phylogenetic diversity (PD), (B) mean pairwise distance (MPD), and (C) mean nearest taxon distance (MNTD) among undisturbed old-growth forest (UO), disturbed old-growth forest (DO), and regrowth forest (R). Hinges represent the 25^th^, 50^th^, and 75^th^ percentiles, respectively. Whiskers extend to maximum 1.5 times the interquartile range. Letters code for significant differences among forest categories.

The regrowth forests, on the other hand, held a higher terminal phylogenetic diversity as compared to both undisturbed and disturbed old-growth forests. Specifically, MNTD was 14.16 % and 17.10 % higher in R than in UO (p < 0.05) and DO (p < 0.05), respectively. For PD and MPD, no significant differences in SES between R and UO were found, nor between R and DO. The phylogenetic structure in R seemed to be more positive in all diversity metrics as compared to UO. However, also here, very few plots showed significant clustering or overdispersion relative to the null model.

### Functional diversity

Overall, the disturbed old-growth forests exhibited a higher functional divergence of traits as compared to undisturbed old-growth forest (Figure 3). Specifically, FDiv was 18.31 % higher in DO (p < 0.01) than in UO. For, FRic, FEve, FDis, and FRaoQ no significant differences were detected among forest categories. In terms of structure of the functional traits based on the SES values, overall DO tended to be more overdispersed than UO, except for FRic, but no plots showed significance relative to the null model.

**Figure 3:**
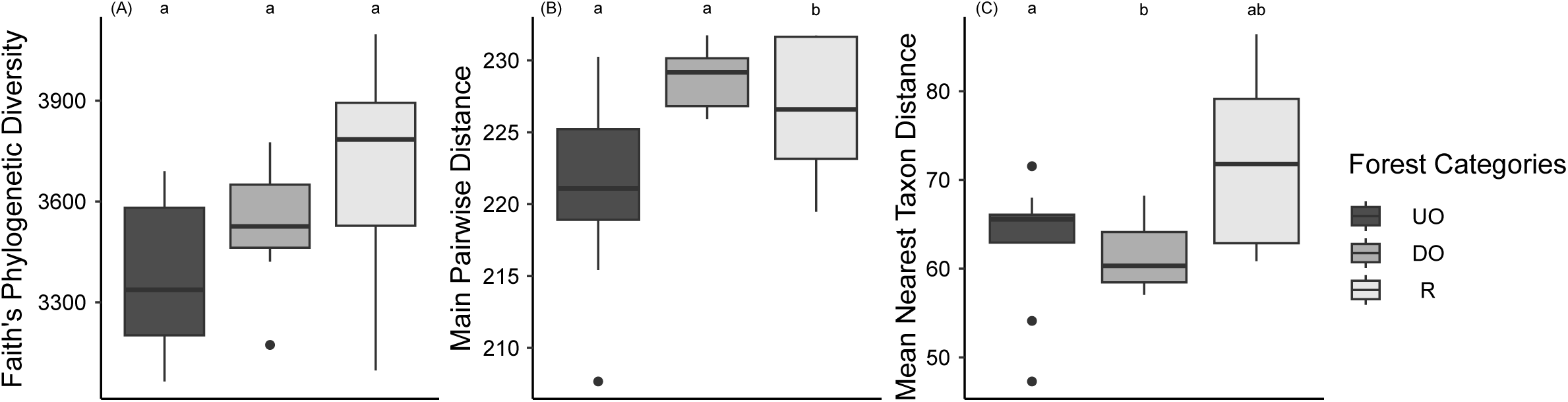
Comparisons of the observed (A) functional richness (Fric), (B) functional divergence (FDiv), (C) functional evenness (FEve), (D) functional dispersion (FDis), and (E) Rao’s quadratic entropy (FRaoQ) among undisturbed old-growth forest (UO), disturbed old-growth forest (DO), and regrowth forest (R). Hinges represent the 25^th^, 50^th^, and 75^th^ percentiles, respectively. Whiskers extend to maximum 1.5 times the interquartile range. Letters code for significant differences among forest categories.

**Figure 4:**
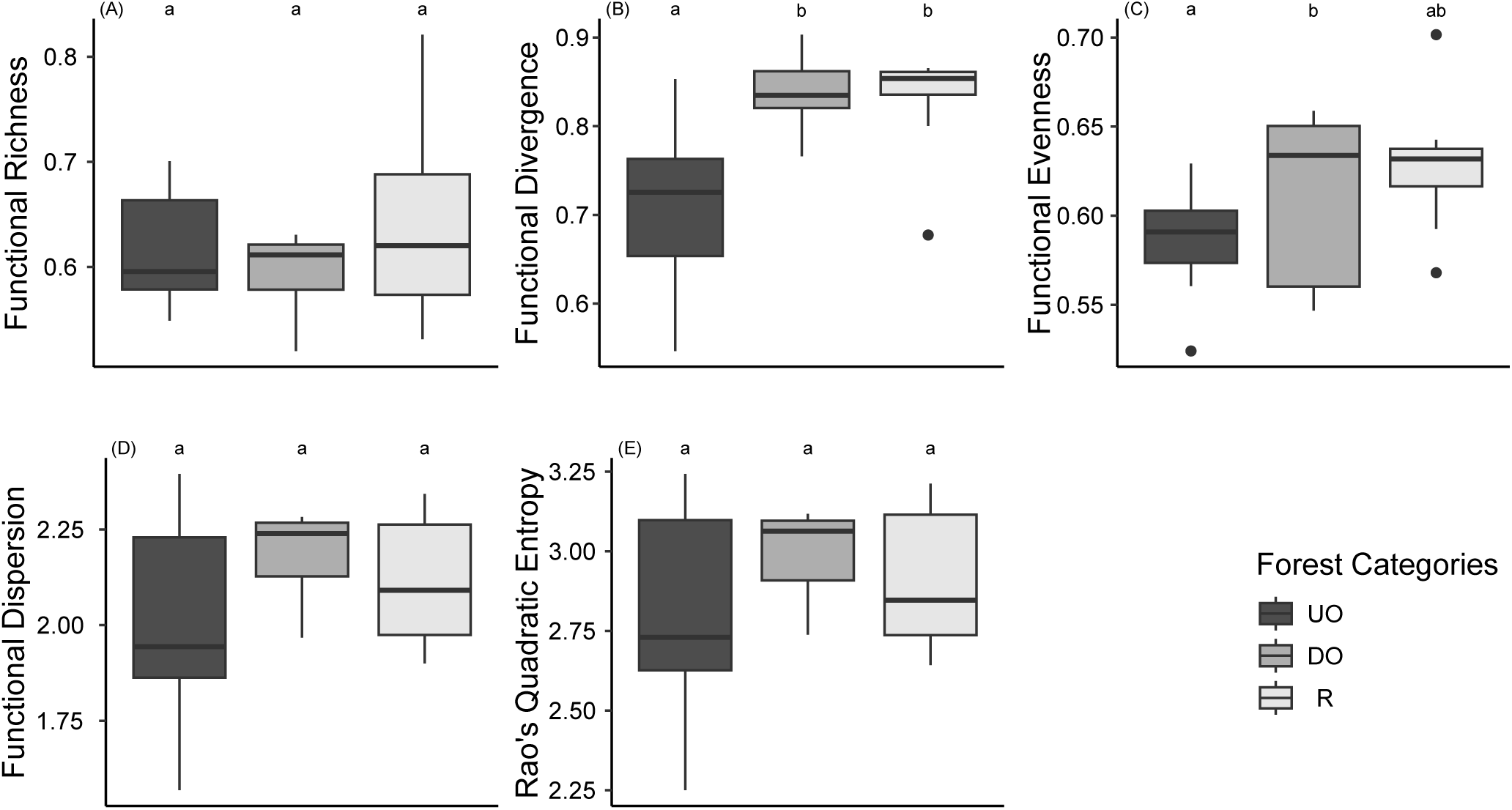
Non-metric multidimensional scaling (NMDS) ordination of the 25 forest inventory plots in the Yangambi region (DR Congo) across three forest categories: undisturbed old-growth forest (UO), disturbed old-growth forest (DO), and regrowth forest (R). Ordinations are based on MNTD (A) phylogenetic dissimilarity and (B) functional dissimilarity. Arrows represent significant drivers of the dissimilarities based on the environmental fitting test analysis (envfit). MAG: magnoliids, ROS: rosids, MON: monocots, FDiv: functional divergence, FEve: functional evenness, WD: wood density, FT1: first fruit type PCOA dimension, FT2: second fruit type PCOA dimension, FT3: third fruit type PCOA dimension, DBH1: first DBH quartile, DBH2: second DBH quartile, DBH3: third DBH quartile, MNTD: mean nearest taxon distance.

A similar trend was observed in regrowth forest, where the functional divergence was higher as compared to undisturbed old-growth forest. Specifically, FDiv was 16.90 % higher in R (p < 0.01) than in UO. The structuring of the functional diversity in R tended to be more overdispersed than UO. Here as well, no plots showed significant clustering or overdispersion when compared to the null model.

### Phylogenetic-functional framework

None of the correlations between phylogenetic and functional diversity metrics were significant (r > 0.6 and p < 0.05), except for MPD and FDiv (r = 0.70; p < 0.001), MPD and FDis (r = 0.68: p < 0.001), and MPD and FRaoq (r = 0.65; p < 0.001) (Table 1). When considering the phylogenetic and functional beta-diversity (dissimilarity among plots), the mantel test did show significant correlations for both phylogenetic and functional dissimilarities. Specifically, a high correlation was found between phylogenetic and functional dissimilarity based on MNTD (r = 0.83; p < 0.001), whereas a moderate correlation was found when both dissimilarities were based on MPD (r = 0.45; p < 0.001) (Table 1). Thus, here we focused on MNTD dissimilarities, while the phylogenetic-functional framework based on MPD dissimilarities is reported in the supplementary material.

**Table 1:** Pearson correlation (r) with corresponding p-values between phylogenetic and functional diversity metrics.

|  | PEARSON CORRELATION | P-VALUE |
| --- | --- | --- |
| PD – FRIC | 0.491 | < 0.05 |
| MPD – FRIC | -0.287 | > 0.05 |
| MNTD – FRIC | -0.120 | > 0.05 |
| PD – FDIV | 0.356 | > 0.05 |
| MPD – FDIV | 0.701 | < 0.001 |
| MNTD – FDIV | 0.180 | > 0.05 |
| PD – FDIS | 0.120 | > 0.05 |
| MPD – FDIS | 0.680 | < 0.001 |
| MNTD – FDIS | 0.060 | > 0.05 |
| PD – FEVE | 0.371 | > 0.05 |
| MPD – FEVE | 0.258 | > 0.05 |
| MNTD – FEVE | 0.421 | < 0.05 |
| PD – FRAOQ | 0.07 | > 0.05 |
| MPD – FRAOQ | 0.654 | < 0.001 |
| MNTD – FRAOQ | 0.041 | > 0.05 |

Both the NMDS based on phylogenetic and functional dissimilarities showed a clear separation of DO forests from UO forests. This visual pattern was further supported by the permutational multivariate analysis of variance which indicated significant differences among forest categories in both phylogenetic (R^2^ = 0.32; p < 0.001) and functional (R^2^ = 0.34; p < 0.001) community composition. Specifically, the DO forests were significantly different from UO in terms of phylogenetic (R^2^ = 0.18; p < 0.01) and functional (R^2^ = 0.21; p < 0.05) community composition.

Likewise, a visual separation of R forests from UO forests was clear in both NMDS ordinations based on phylogenetic and functional dissimilarities. Again, the permutational multivariate analysis of variance supported this visual pattern and indicated significant differences between R forest and UO in terms of phylogenetic (R^2^ = 0.38; p < 0.001) and functional (R^2^ = 0.41; p < 0.001) community composition.

The phylogenetic lineages contributing to the observed patterns of phylogenetic dissimilarity were the Magnoliids (R² = 0.03; p < 0.05) and the Rosids (R² = 0.55; p < 0.001). Similarly, for functional dissimilarity, the Magnoliids (R² = 0.34; p < 0.05) and Rosids (R² = 0.57; p < 0.001) were significant drivers, with the Monocots (R² = 0.27; p < 0.05) also showing a significant contribution. Furthermore, drivers of changes in phylogenetic and functional composition among the different forest categories are reported in table 2.

**Table 2:** Drivers of the changes in phylogenetic and functional composition, with corresponding R^2^ and p-values.

|  | PHYLOGENETIC COMPOSITION |  | FUNCTIONAL COMPOSITION |  |
| --- | --- | --- | --- | --- |
|  | **R<sup>2</sup>** | **p-value** | **R<sup>2</sup>** | **p-value** |
| **FDIV** | 0.46 | < 0.01 | 0.41 | < 0.01 |
| **DBH1** | 0.31 | < 0.05 | 0.31 | < 0.05 |
| **DBH2** | 0.42 | < 0.01 | 0.43 | < 0.01 |
| **DBH3** | 0.34 | < 0.05 | 0.43 | < 0.01 |
| **WD** | 0.7 | < 0.01 | 0.62 | < 0.01 |
| **FT1** | 0.39 | < 0.01 | 0.48 | < 0.01 |
| **FT2** | 0.68 | < 0.01 | 0.58 | < 0.01 |
| **FT3** | 0.33 | < 0.05 | 0.30 | < 0.01 |
| **FEVE** | - | - | 0.40 | < 0.01 |
| **MNTD** | - | - | 0.28 | < 0.01 |

## Discussion

Assessing the phylogenetic and functional dimensions is key to the effective conservation of globally important ecosystems like the tropical rainforests. Here, by integrating these dimensions with forest inventories of Depecker *et al*. (2022), we were able to quantify changes in phylogenetic and functional community composition driven by changes in trunk size, fruit type, and mainly wood density, all of which are associated with selective logging and forest regrowth. These changes can affect the strength of ecosystem functions and may pose significant risks to the stability of tropical rainforests in the Congo Basin.

### Selective logging

The effects of selective logging on the different diversity measures were assessed by comparing disturbed old-growth forests with undisturbed old-growth forests. The most prominent observation was that selective logging was clearly associated with alterations in the phylogenetic composition paralleled by changes in the functional trait composition, whereas a less noticeable impact on the phylogenetic and functional diversity was observed. This result was in line with our hypothesis. Indeed, in the Yangambi area, exploitable tree species are selectively removed from the forest, strongly reducing the number of individuals but not the number of species (Depecker *et al*., 2022). Tree species that are commonly targeted by loggers are *Prioria oxyphylla* (Harms) Breteler and *Strombosia pustulata* Oliv., which are not deeply rooted in the phylogenetic tree. The reduced dominance of these species might explain the slight increase in phylogenetic diversity of more basally rooted lineages in the phylogenetic tree in disturbed old-growth forests. Additionally, the phylogenetic structure indicates that the effects of selective logging are relatively limited, as there is no sudden shift towards species being more clustered or overdispersed (Letcher, 2010). This could be related to the fact that logging primarily impacts the number of individuals from species that are evenly distributed throughout the phylogenetic tree, suggesting that species or lineages are not entirely lost, as we hypothesised. Likewise, Arroyo-Rodríguez *et al*. (2012) reported few changes in phylogenetic diversity and structure in South American tropical rainforests despite high levels of forest disturbance, and similar results were reported elsewhere in the Neotropics (Matos *et al*., 2017; Santo-Silva *et al*., 2018; Rocha-Santos *et al*., 2023). These studies additionally concluded that disturbed forests still hold important conservation value and can likely sustain forest ecosystem functioning (Cadotte *et al*., 2012; Santo-Silva *et al*., 2018; Rocha-Santos *et al*., 2023). Our results on functional diversity confirm this conclusion, as no major changes associated with disturbance were reported in any functional diversity metric, except for functional divergence which reflects how abundance is spread along the different traits (Grenié & Gruson, 2023) and which is likely a direct effect of the reduction in the dominance of certain species. Similar results were reported in tropical rainforests in Bolivia and French Guiana (Baraloto *et al*., 2012; Carreño-Rocabado *et al*., 2012). It is, however, noteworthy that phylogenetic diversity was in general not correlated with functional diversity, and which was also the conclusion from a study in Brazil (Rocha-Santos *et al*., 2023), but in sharp contrast with the relationship between phylogenetic and functional diversity reported in literature (Srivastava *et al*., 2012; Tucker *et al*., 2018). Our study hereby contributes to the evidence emphasising the importance of using both phylogenetic and functional diversity in tandem (Devictor *et al*., 2010; Mazel *et al*., 2018; Tucker *et al*., 2018).

Alpha-diversity measures are, however, just one facet of the total diversity spectrum (Chave *et al*., 2007; Hernández□Ordóñez *et al*., 2019). The phylogenetic and functional composition (cf. beta-diversity or dissimilarity) is crucial to evaluate subtle effects of selective logging and is furthermore key to connecting local to regional processes (Santo-Silva *et al*., 2018). Our results confirm that selective logging has resulted in a significantly altered community composition in both phylogenetic and functional dimensions, and that these dimensions are highly correlated. This observation is corroborated by studies on forests in other tropical regions (Baraloto *et al*., 2012; Carreño-Rocabado *et al*., 2012; Ding *et al*., 2019). In terms of the phylogenetic dimension, the dissimilarity between the different forest inventory plots is mainly driven by changes in the composition of Rosids, the clade to which the most targeted tree species by logging belong, like *P. oxyphylla.* This is, however, not surprising as the Rosids are one of the most diversified clades globally (Zuntini *et al*., 2024), especially in the tropics, where the clade is ecologically dominant and has its highest species richness, with numerous species with relatively high wood density (Chave *et al*., 2006; Swenson & Enquist, 2007; Folk *et al*., 2018; Sun *et al*., 2020).

The phylogenetic and functional dissimilarity in tree communities is driven by a lower wood density and larger-sized trunk widths associated with selective logging. The trees in the Yangambi area that are selectively removed are mainly species with a high wood density, as these are most suitable for construction material and the production of charcoal (Larjavaara & Muller-Landau, 2010; Couto *et al*., 2023). Studies with indigenous human communities in the western Amazon region (De Aledo *et al*., 2023), in human modified forests in the eastern Amazon region (Berenguer *et al*., 2018), and along a forest disturbance gradient in Madagascar (Brown *et al*., 2013), all confirm the preference for highly dense wood for construction and production of charcoal, and further confirm the resulting impact on the remaining forests. As the logging in Yangambi is a purely manual labour this likely explains the preference for logging smaller sized trees, leading to a shift towards larger-size trunk widths in disturbed old-growth forests. In addition to the community compositional changes in wood density and trunk width, our study also revealed that selective logging can be associated with changes in fruit types. Specifically, less berry and drupe harbouring trees are to be found in disturbed old-growth forests as compared to undisturbed old-growth forests. This trend towards less fleshy fruits is like to be an indirect effect rather than a direct effect of selective logging. Yet, including fruit type in studies on the effects of selective logging is crucial, as it can facilitate future monitoring of changes in species composition, including animal populations contributing to fruit dispersal. Furthermore, the shift towards a generally lower wood density in communities in disturbed old-growth forests is likely also affecting their capacity for carbon storage, as trees with a low wood density are not able to sequester and store carbon as efficiently as trees with a higher wood density (Chave *et al*., 2005; Chave *et al*., 2009). Thus, although the ecosystem functions of the rainforests here seem to be conserved, or at the very least not reduced in terms of functional diversity, the strength of the provided ecosystem functions is very likely to be altered by the changes in the community compositions. This could be particularly important, especially concerning the trees’ reproductive traits, which could affect faunal composition and dispersal.

### Forest regrowth following agricultural abandonment

The outcomes of forest regrowth were assessed by comparing undisturbed old-growth forests with regrowth forests on agricultural lands which were abandoned somewhere between 40 and 60 years ago. Like in undisturbed old-growth forests, no significant clustering, nor overdispersion of phylogenetic lineages and functional traits was detected. Similar to studies along successional gradients in China (Mo *et al*., 2013), Borneo (Mahayani *et al*., 2020; Mahayani *et al*., 2022), and Costa Rica (Letcher, 2010), this would indicate that the forests were able to recover in terms of phylogenetic and functional structure. This also holds true for the functional dimension of the regrowth forests, which is entirely similar to the functional diversity recorded in undisturbed old-growth forests, except for the functional divergence. These results are in line with the few other studies on this topic in the Congo Basin (Bauters *et al*., 2019; Makelele *et al*., 2021). The phylogenetic diversity, on the other hand, seemed not fully recovered to levels comparable to undisturbed old-growth forest. Tree species like *Myrianthus arboreus* P. Beauv. and *Dacryodes edulis* G. Don, a late pioneer species and a secondary forest species, respectively, were characteristic for the regrowth forests (Meunier *et al*., 2015; PROTA, 2021; Depecker *et al*., 2022) and are rooted more terminally in the phylogenetic tree, possibly explaining the higher terminal phylogenetic diversity in these forests. The presence of these trees combined with the higher phylogenetic diversity support the idea that the regrowth forests are in a late transitional state towards an old-growth forest, but not yet fully recovered. The phylogenetic and functional composition in the regrowth forests seems to confirm this point of view, as these were still significantly different from compositions recorded in the undisturbed old-growth forests. Early-successional species have a lifespan of around 30 years, whereas late-successional species can last up to 150 years (Mo *et al*., 2013; DeArmond *et al*., 2022). This longer lifespan exceeds the time that the forest has had to regrow, which likely explains why late-successional species are still present in the communities of regrowth forests. These late-successional-species commonly have lower wood density and typically harbour dry fruits like capsules and achenes (Larjavaara & Muller-Landau, 2010), which are among the most important drivers of the dissimilarities between regrowth forests and undisturbed old-growth forests. Surprisingly, trunk width also contributes to the difference between regrowth forests and undisturbed old-growth forests, but in an unexpected opposite way: undisturbed old-growth forests had a greater number of trees with small trunks. A similar pattern was observed in disturbed old-growth forests, suggesting that anthropogenic activities are likely hampering the recovery of regrowth forests.

### Limitations

As for the majority of studies in the Afrotropics, our study was limited by the availability of both phylogenetic and trait data (Erickson *et al*., 2014). Of the 211 sampled taxa in Depecker *et al*. (2022), only 167 were identified at species level, potentially leading to a less accurate estimate of phylogenetic diversity (Tucker & Cadotte, 2013). However, this issue is consistent across all forest categories and has, therefore, limited influence on the outcome of this comparative study. Additionally, the correlation between the number of species and phylogenetic diversity was accounted for by calculating standardised effect sizes of the corresponding metrics (Mazel *et al*., 2016). Likewise, functional diversity depends on the *a priori* choice of traits (Baraloto *et al*., 2012). Although limited, the traits included in this study were chosen to represent usefulness, vegetative, and reproductive functions, as it was reported that excessive incorporation of correlated traits can result in biased estimates of functional diversity and only traits related to the functions of interest should be included (Cadotte *et al*., 2011). A final limitation, inherent to studies in tropical forests, is the spatial clustering of the forest inventory plots (Wittmann *et al*., 2006; Liebsch *et al*., 2008; Ganivet *et al*., 2020). This has been accounted for as much as possible in the analyses and interpretation.

## Conclusion

Our study shows that incorporating phylogenetic and functional dimensions of diversity in assessing the effects of selective logging and forest regrowth on tropical African rainforests can yield novel insights in tree community dynamics resulting from anthropogenic disturbance. We found significantly altered phylogenetic and functional compositions in disturbed old-growth forests and regrowth forests, as compared to reference undisturbed old-growth forest. The dissimilarities of the communities were driven by changes in trunk size and fruit type, but mainly by changes in wood density, which was clearly associated with selective logging and forest regrowth. The phylogenetic and functional diversity, on the other hand, seemed to be less affected by selective logging and has nearly recovered 40 to 60 years after agricultural abandonment. Our results indicate that the ecosystem functions are maintained in the disturbed old-growth forests and recovered in the regrowth forests, but the strength of the functions is severely reduced. The conservation of undisturbed old-growth forests is thus key in this everchanging world, as they have been proven irreplaceable (Gibson *et al*., 2011). These old-growth forests can, furthermore, be complemented by regrowth forests as they contribute relatively fast to recovering biodiversity and ecosystem functions.

## Acknowledgements

We would like to thank the Institut National pour l’Etude et la Recherche Agronomiques (INERA) and the FORETS project, which is financed by the 11th European Development Fund, and the University of Pisa for facilitating this study.

## Author contributions

JD and FV conceived this study, and AC and JD further designed it. JD analysed the data. JD, FV, OH, SJ, and AC wrote the manuscript. All authors contributed to finalising the manuscript

## Data availability statement

The data that support the findings of this study will be made openly available on Zenodo.

## Supplementary material

**Table S1:** Trait values per species. WD: wood density; SLA: specific leaf area: FT: fruit type; WD ref: source of wood density data; SLA ref: source of specific leaf area data, herbarium specimen used for measurements; FT ref: source of fruit type data; PT ref: source of assessment of the species’ closest relative.

**Figure S1:** Comparisons of the standardised effect sizes (SES) of (A) Faith’s phylogenetic diversity (PD), (B) mean pairwise distance (MPD), and (C) mean nearest taxon distance (MNTD) among undisturbed old-growth forest (UO), disturbed old-growth forest (DO), and regrowth forest (R). Hinges represent the 25^th^, 50^th^, and 75^th^ percentiles, respectively. Whiskers extend to maximum 1.5 times the interquartile range.

**Figure S2:** Comparisons of the standardised effect sizes (SES) of (A) functional richness (Fric), (B) functional divergence (FDiv), (C) functional evenness (FEve), (D) functional dispersion (FDis), and (E) Rao’s quadratic entropy (FRaoQ) among undisturbed old-growth forest (UO), disturbed old-growth forest (DO), and regrowth forest (R). Hinges represent the 25^th^, 50^th^, and 75^th^ percentiles, respectively. Whiskers extend to maximum 1.5 times the interquartile range.

**Figure S3:** Non-metric multidimensional scaling (NMDS) ordination of the 25 forest inventory plots in the Yangambi region (DR Congo) across three forest categories: undisturbed old-growth forest (UO), disturbed old-growth forest (DO), and regrowth forest (R). Ordinations are based on MPD (A) phylogenetic dissimilarity and (B) functional dissimilarity. Arrows represent significant drivers of the dissimilarities based on the environmental fitting test analysis (envfit). FDiv: functional divergence, FEve: functional evenness, WD: wood density, FT1: first fruit type PCOA dimension, FT2: second fruit type PCOA dimension, FT3: third fruit type PCOA dimension, DBH2: second DBH quartile, PD: Faith’s phylogenetic diversity.

