## Supplementary figures and images for "Phylogenetic and functional compositional shifts associated with selective logging and forest regrowth in tropical rainforests of the Congo Basin"

### Supplemental figure S1

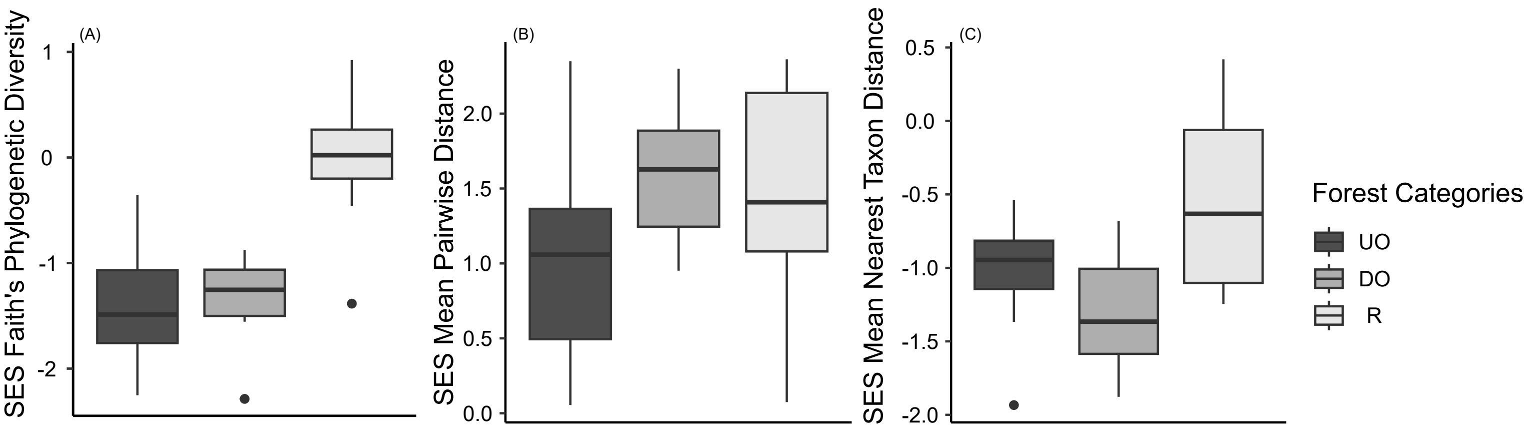

### Supplemental figure S2

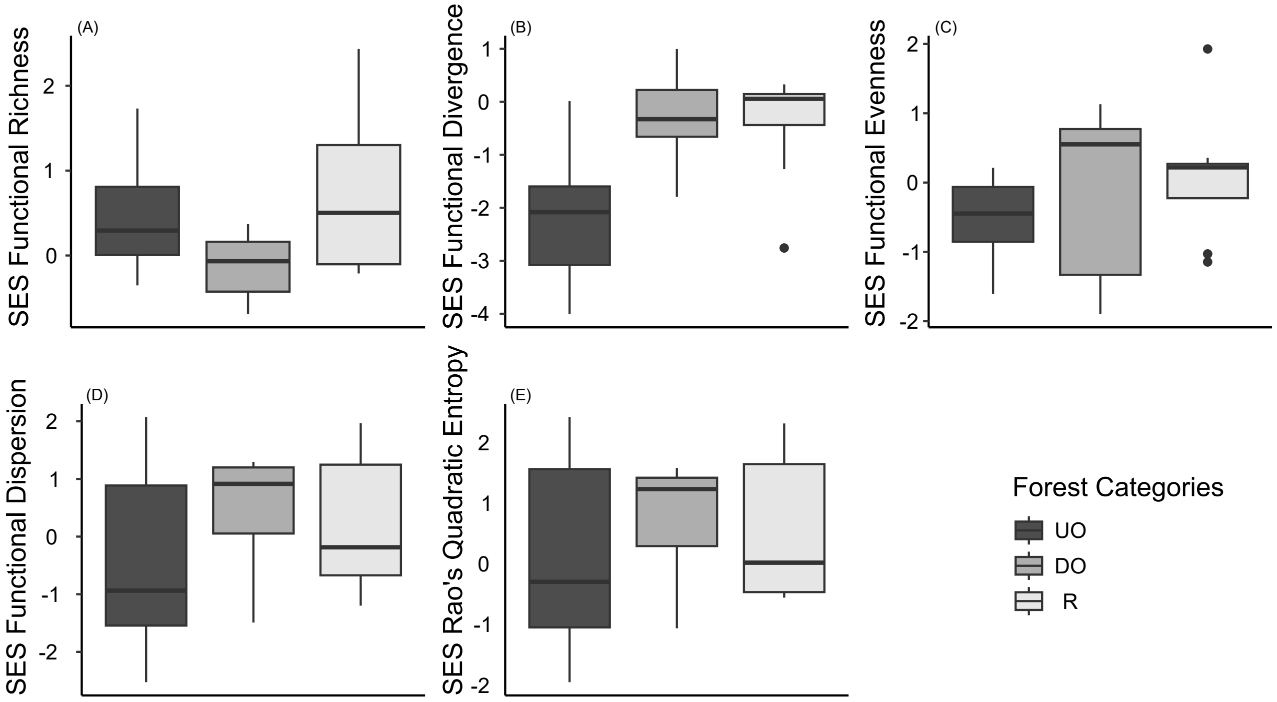

### Supplemental figure S3

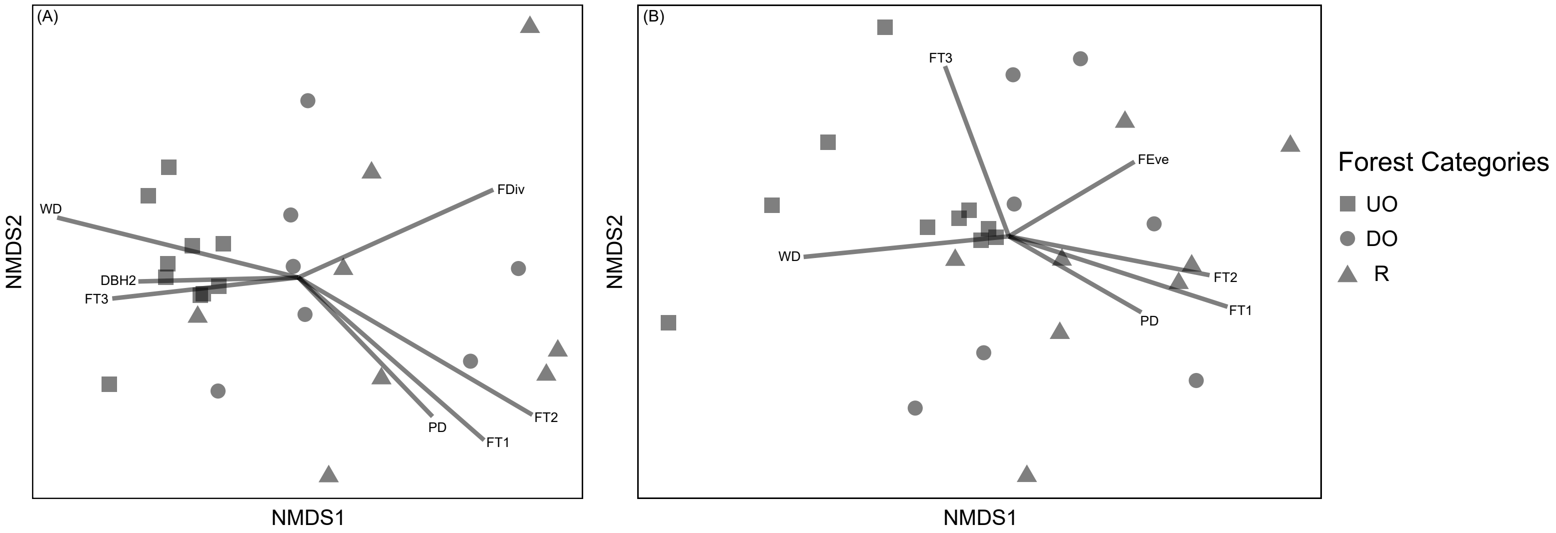
